# REFinder: Mining new enzymes from metagenomes

**DOI:** 10.64898/2026.10.01.756075

**Authors:** R. Prabakaran, Yana Bromberg

**Affiliations:** Department of Biology, Emory University, Atlanta, GA 30322, USA; Department of Computer Science, Emory University, Atlanta, GA 30322, USA

**Keywords:** Functional Metagenomics, Enzymes, Catalytic activity, Protein Function, Pipeline, Bioinformatics, Language model

## Abstract

Microbial metagenomes encode vast catalytic diversity, but recovering enzymes of novel, yet-undescribed functionality typically requires whole-community assembly plus a means of identifying active proteins *ab initio*. The first step of the task is compute-intensive and misses low-abundance sequences. The second is complicated by our inability to predict novel functions. REBEAN, our DNA language model, sidesteps the latter by assigning each sequencing read a high-level Enzyme Commission (EC) class or a non-enzyme label without alignment. Here we build REFinder, a pipeline that addresses both steps by routing REBEAN-annotated reads to assemble only the putative enzymatic reads that have identified catalytic signatures.

In our evaluation of 50 microbiome metagenomes, REFinder identified 1.2 to 2.1 fold more enzymes than could be recovered via homology-based annotation of the proteins from the corresponding full assemblies. Moreover, it was as much as 6.4-fold cheaper computationally than full assembly. Across all samples, REFinder identified at least three fourths and as many as 90% of the homology-accessible enzymes identified via full assembly of the complete metagenomes. Notably, a fraction of these, 22% to 45% per EC class, carried no similarity to Swiss-Prot proteins, i.e. a set of enzymes homology cannot annotate. For roughly two fifths of the over thirteen thousand such novel oxidoreductases from two saliva samples, ESMFold predicted structures aligned with TM-score≥0.7 to a characterized enzyme structure in the PDB – a substantial structural similarity without sequence homology. These results illustrate that targeted, alignment-free assembly turns even well-mapped microbiomes into a source of thousands of previously invisible but credible novel enzymes, raising our expectations for exploration of environmental microbiomes.

**Availability and implementation:** https://bitbucket.org/bromberglab/NF-REFinder/

## 1. Introduction

The majority of environmental microbial taxa resist standard laboratory cultivation ^1^. Thus, metagenome sequencing, i.e. sequencing an environmental sample as a whole, is the most common path to identifying genes/proteins encoded in these unculturable lineages ^2^. Metagenome mining enables identification of novel biocatalysts, providing enzymes with enhanced thermal stability, activity profiles, and, potentially, even new catalytic mechanisms ^3-5^.

How do we identify novel functionality encoded in metagenomic samples? Two routes are salient – (1) assembling metagenomes to long contigs/MAGs (metagenome assembled genomes) and finding/labeling encoded genes or (2) annotating functionality directly from reads. Existing read-level annotators, e.g. MG-RAST ^6^ and mi-faser ^7^, label individual reads by molecular function or enzyme class via calibrated homology to known genes. Read-based profilers such as HUMAnN quantify pathways and gene families in a sample ^8^. These approaches are relatively straightforward, but since a short read spans only a fragment of a gene, read-level homology annotation is less complete and less reliable than full-length-gene annotation ^9^ and often cannot resolve the specific gene of read-origin or its function ^10^. The field as a whole, therefore, deems the first route, i.e. assemble-then-annotate, where annotation runs on longer, near-complete gene (or translated protein) sequences ^11, 12^ more reliable.

Read- and assembled gene-based annotations share a constraint in that both assign function by aligning sequences to reference databases such as UniProt, KEGG, and/or eggNOG ^13, 14^. Multiple algorithms for alignment and scoring alignments in light of functional similarity have been developed ^7, 15, 16^. Recent advances in structure prediction have also facilitated functional annotation via structural alignments ^17, 18^. In either case, however, alignment can only recognize that which resembles a known reference. Novel or divergent enzymes, i.e. the functional dark matter that dominates many microbiomes, are therefore missed by default, whether annotation runs on reads or on contigs ^19^.

We previously developed REBEAN – a model that removes the homology need at the read level. Built as a DNA language model, it assigns each read an Enzyme Commission class (EC 1–7, ^20^) or a non-enzyme label from sequence context alone ^21^. Because it reads functional signal from patterns rather than reference matches, and with read embeddings largely independent of sequence identity ^21^, REBEAN recovers enzymatic reads that alignment overlooks: three-to six-fold more enzymatic reads. We could thus choose which reads to assemble (**Fig. 1**), routing only those labeled with a target EC class and yielding an activity-focused assembly (seeding) at a fraction of the cost of assembling everything, i.e. avoiding the step that dominates metagenomic compute ^22, 23^. An idea similar to targeted assembly of specific protein families that has been demonstrated to accelerate discovery ^23-27^, but carried out through a DNA language model, targeting functional activities.

**Figure 1:**
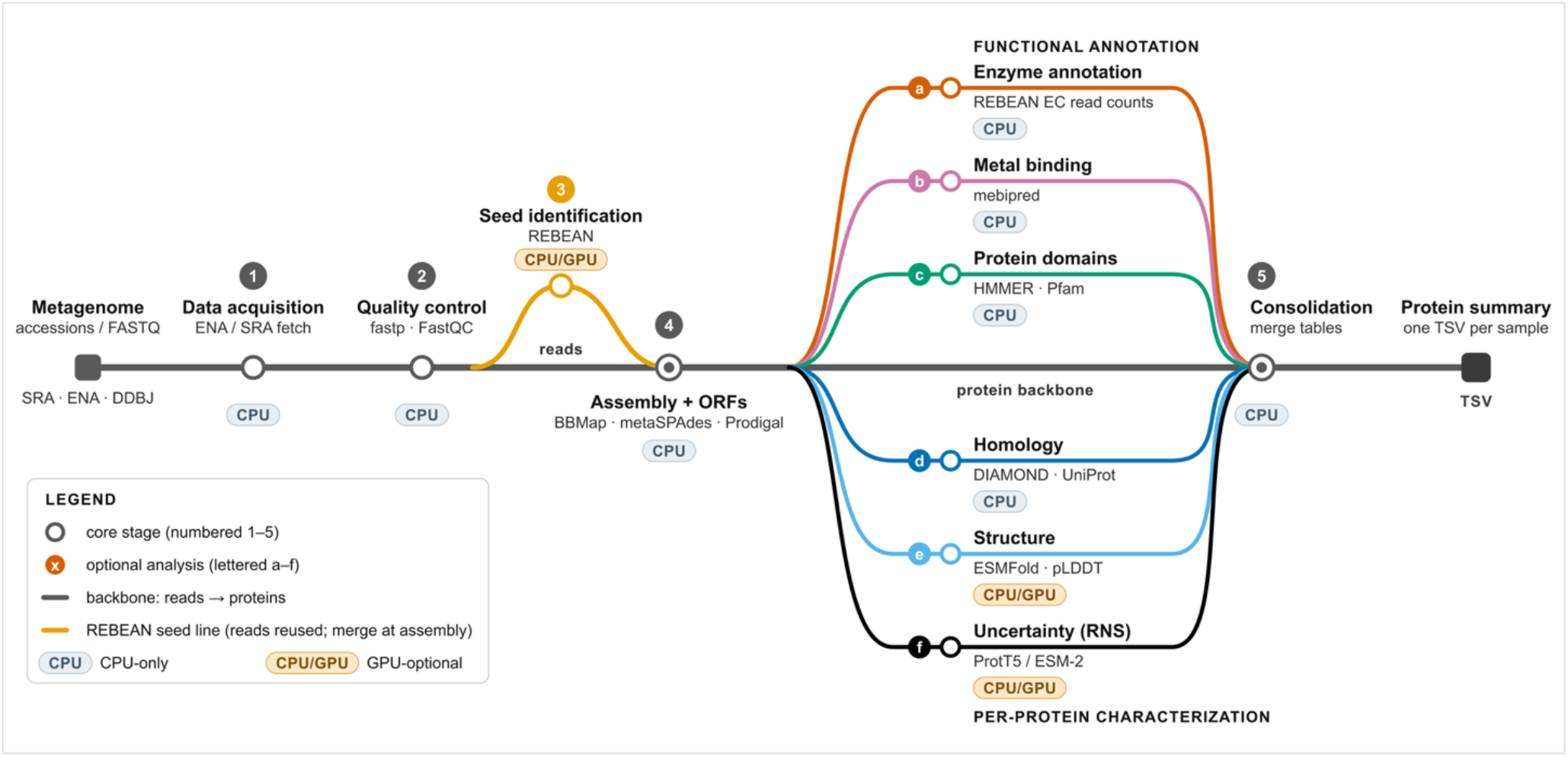
The REFinder pipeline. A Nextflow workflow that turns raw shotgun-metagenome reads into a functionally annotated protein table. *Inputs:* sample accessions (ENA/SRA). *Core stages:* (1) reads are fetched from source repositories, (2) quality-trimmed (fastp, FastQC); (3) REBEAN classifies every read into EC1–7 or non-enzyme, yielding per-class seeds; (4) the reads making up each seed are recruited with BBMap (minid 0.90), assembled with metaSPAdes (– meta), and gene-called with Prodigal; (5) outputs are consolidated into a protein table. *Optional annotation modules* include: DIAMOND homology (EC numbers), HMMER against Pfam-A, ESMFold structure prediction, mebiPred metal-binding prediction, and ESM-2/ProtT5 embedding reliability (RNS) scoring.

In this work, we use 50 metagenomic samples from two studies to evaluate our seeding pipeline’s ability to recover enzymes vs. a full assembly-driven pipeline. Our results highlight the potential to reduce computational costs of metagenomic analyses while expanding our understanding of biological catalytic mechanisms.

## 2. Methods

### 2.1. The REFinder pipeline

We built REBEAN_Enzyme_Finder (REFinder), a read seed-based enzyme-discovery pipeline (**Fig. 1**). REFinder is a Nextflow ^28^ read-to-annotation workflow built around REBEAN ^21^, a DNA language model-based classifier that labels each sequencing read into one of seven Enzyme Commission classes (EC1–7, ^20^) or as non-enzyme. REFinder has five core stages and six optional annotation modules. The core stages are:

1. Data acquisition fetching reads from ENA/SRA
2. Quality Control (QC) - quality-trimming with fastp ^29^ and FastQC ^30^.
3. Enzyme-read detection, in which REBEAN scores every read; selected reads of each class form that class’s seed.
4. Seed-guided assembly, in which i) BBMap ^31^ recruits each seed’s read neighborhood from the complete sample (minid 0.90), ii) metaSPAdes (–meta; ^11^) assembles the recruited reads, and iii) Prodigal (-p meta; ^12^) predicts genes and translates them into proteins.
5. Consolidation of all outputs into one per-protein table.

The six optional protein-annotation modules are:

a. Enzyme annotation of proteins through mapping of REBEAN identified enzymatic reads
b. Metal-binding prediction with MebiPred ^32^
c. Homology search against Pfam-A families ^33^ through HMM search using HMMER ^34^
d. Homology search against Swiss-Prot and/or TrEMBL ^35^ through alignment using DIAMOND ^36^ to assign EC numbers.
e. Protein structure prediction using ESMFold v1^37^
f. Reliability estimation ^38^ of protein embedding, with an option to choose between two commonly used protein language models: ESM-2 (esm2_t33_650M_UR50D) ProtT5 (prot_t5_xl_half_uniref50-enc) ^37, 39^ .

#### Data

We evaluated REFinder on two publicly available metagenome sets: human saliva microbiome (BioProject PRJNA396840 ^40^, *n* = 30) and murine antibiotic-perturbation of the gut microbiome study (BioProject PRJNA504846 ^41^, *n* = 20). Note that we chose these easily accessible and well-studied environments deliberately as a conservative test for evaluation of our method vs. a traditional full-assembly homology-based pipeline. Human and mouse-associated microbiomes are less-diverse than most environmental samples and are among the most deeply sequenced microbiomes available. Thus, any enzyme that REFinder surfaces here is unlikely to be an artifact of shallow sampling. However, even in this well-mapped setting a large fraction of enzymes per class had no close Swiss-Prot matches (**Section 3.2**).

As per our workflow, metagenomic reads were quality-trimmed with fastp (-q 20, –length_required 50, adapter auto-detection; ^29^) resulting in a mean read length of 244.9 ± 13.8 bp (saliva samples) and 149.4 ± 7.4 bp (gut samples), and then processed through two routes: (1) a full community assembly (full arm), and (2) seven REFinder runs, one per EC class (EC-*N* = 1 ™ 7 seed arm). The full arm assembled every trimmed read. Each seed arm, instead, assembled one EC class; i.e. it took as seeds all reads that REBEAN labeled with that class, recruiting their read neighborhood from the full sample with BBMap (minid 0.90, ^31^). For both full-assembly and seeded routes, reads were assembled with metaSPAdes 4.2.0 (–meta, *k* = 21,33,55; ^11^), and gene-called/translated into protein sequences with Prodigal 2.6.3 (-p meta; ^12^).

### 2.2. Annotation of genes as enzymes

Predicted proteins from both assembly arms were annotated as enzymes in two independent ways:

#### 2.2.1. Homology-based annotation

A predicted protein was deemed a true enzyme if and only if its best Swiss-Prot hit ^35^, assigned with DIAMOND 2.1.12 blastp (best hit, e-value ≤1E-5; ^36^), carried exactly one EC number. Note that we performed function annotation transfer without a minimum identity or coverage threshold to maximize the recovery instead of relying on quality-centric, but low recall, approaches (e.g. ^15^). The assigned true EC class was the first digit of that EC number, regardless of its Swiss-Prot evidence code. In addition, each read inherits the Swiss-Prot accession number of the protein’s hit. We used Swiss-Prot, the manually curated subset of UniProt as reference, rather than the automatically annotated TrEMBL whose annotations are derived from a Swiss-Prot subset. This is computationally faster while retaining identical coverage of the function space. Note that Swiss-Prot is REFinder’s default when sequence similarity/homology identification is requested, with TrEMBL available as an option.

#### 2.2.2. REBEAN-based definition

A protein was deemed an EC-N enzyme if at least one mapped read’s score reached the EC-N class-specific 90%-precision cutoff (EC1 0.944, EC2 0.971, EC3 0.947, EC4 0.829, EC5 0.888, EC6 0.863, EC7 0.698). Because classes are considered separately, proteins with reads mapping to different EC-N classes were counted multiple times. Unlike the homology-based annotations, this path can identify enzymes with no Swiss-Prot matches.

To create protein-linked identifiers, we clustered all assembly-extracted proteins (per sample, full and all seed arms separately) with MMseqs2 at 90% identity and 80% coverage ^42^. Each cluster was given an identifier, and we then provided each read with the cluster identifier of a protein that the read had mapped to.

### 2.3. Evaluation criteria

We compared REFinder’s REBEAN-seeded assembly enzyme discovery pipeline (seeded arm) against a traditional full-assembly pipeline with homology-based annotation (full arm) on both saliva and murine samples. The comparison was done in two phases: i) evaluating recovery of known enzymes and ii) evaluating the identified novel enzymes. We also compared the computational cost of assembly in each arm.

#### 2.3.1. Recovery

Comparison of known enzyme recovery between the REFinder’s seeded arm and the class-matched full-assembly (full arm) was summarized by overlap of enzymes (**Eq. 1**), where A and B denote i) the distinct Swiss-Prot accession sets identified (**Section 2.2.1**) or ii) number of distinct enzyme clusters (> 90% identity and 80% coverage) (**Section 2.2.2**) assembled, from each arm computed within one EC-N class and one sample.

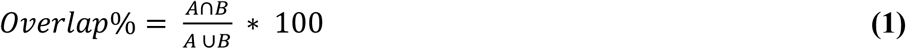

#### 2.3.2. Novel enzymes

A protein labelled as an enzyme via annotation transfer from a Swiss-Prot match (**Section 2.2.1**) was considered novel if the match was <30% amino-acid identity ^36^, i.e. not a reliable call ^43^. Similarly, a protein labelled as an enzyme via REBEAN assigned read labels (**Section 2.2.2**) was considered novel if it had no Swiss-Prot hit or if its best hit fell below 30% sequence identity.

#### 2.3.3. Computational cost

Compute was recorded as CPU-hours for fastp, seed derivation, BBMap recruitment, metaSPAdes, Prodigal, and DIAMOND, summed over the samples of each dataset, and captured with /usr/bin/time -v and slurm accounting (sacct).

## 3. Results and Discussion

### 3.1. Seeded assembly recovers known enzymes

We compared the genes assembled through full-assembly arm and the seeded arm based on their ability to reconstitute homologs of Swiss-Prot enzymes. As expected due to seeding with a prediction model whose recall is below 100% (80.6% for saliva and 84.2% for murine), the seeded arm recovered somewhat fewer (∼76%) homology-identifiable enzymes than the full arm (**Tables 1, S1**).

**Table 1:** Recovery of known enzymes from the saliva metagenomes.

| EC | Class | No. of distinct Swiss-Prot Enzymes identified |  |  |  |  | Overlap % |
| --- | --- | --- | --- | --- | --- | --- | --- |
|  |  | Full arm | <sup>†</sup>Full arm w REBEAN filter | Seeded arm | Unique to full arm | Unique to seeded arm |  |
| 1.* | Oxidoreductase | 2,685±451 | 2,341±392 | 2,381±403 | 126±31 | 166±45 | 88.5±1.3 |
| 2.* | Transferase | 8,082±1462 | 7,132±1,308 | 7,242±1347 | 354±83 | 464±125 | 89.3±1.0 |
| 3.* | Hydrolase | 5,433±931 | 3,850±708 | 3,985±749 | 367±75 | 503±116 | 80.1±1.2 |
| 4.* | Lyase | 1,850±343 | 1,363±255 | 1,420±269 | 102±26 | 159±46 | 83.0±2.0 |
| 5.* | Isomerase | 1,649±295 | 1,279±234 | 1,322±243 | 91±21 | 134±34 | 84.1±1.7 |
| 6.* | Ligase | 3,114±625 | 2,380±469 | 2,518±511 | 190±45 | 328±105 | 81.1±2.4 |
| 7.* | Translocase | 1,522±262 | 1,268±222 | 1,322±233 | 89±19 | 143±37 | 83.7±2.0 |

<sup>†</sup> To ensure a standardized efficiency comparison between the Seeded (REFinder) and Full arm assemblies, we restricted the analysis to genes containing at least one REBEAN-predicted enzymatic read. The mean and standard deviation over the 30 samples (PRJNA396840) is reported. Note counts are number of distinct Swiss-Prot accessions (**Section 2.3.1**).

To isolate the assembly stage from the seeding stage, we evaluated the seeded arm against the subset of full-assembly arm enzymes that REBEAN could detect. We required that, for each gene identified in the full arm, at least one mapped read be predicted to be of the matching EC-*N* class. On this common ground, the two arms were more closely aligned (∼84%), with the seeded arm recovering slightly more enzymes than the full arm in every EC class of the saliva set (n=30) and in five of seven EC classes of the murine gut set (n=20). These results show that the two pipelines can thus be deemed comparable in recovery of known enzymes (**Tables 1, S1 & Figure 2a, S1a**).

**Figure 2:**
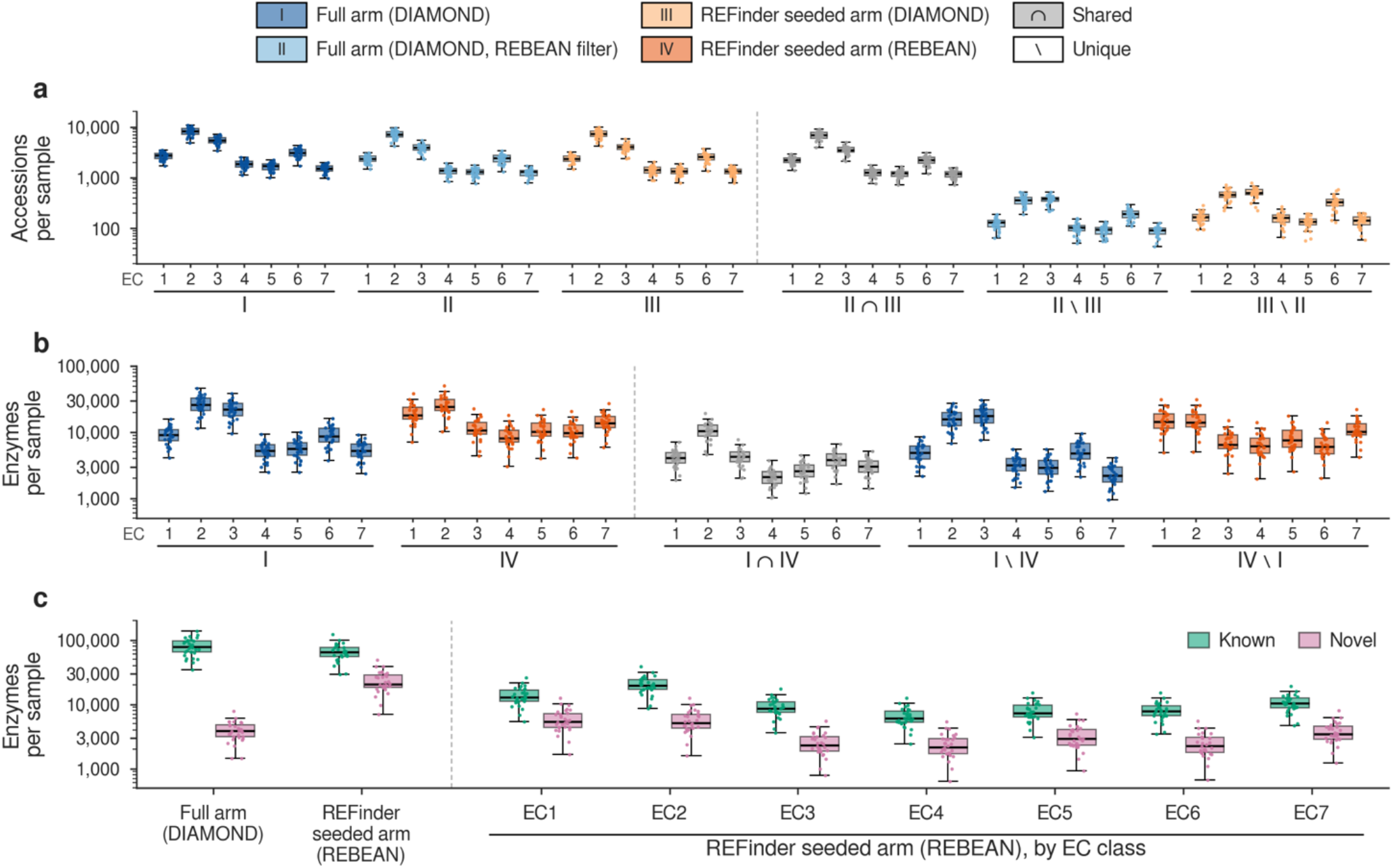
Enzymes recovered from oral saliva metagenomes by full arm and REFinder’s seeded arm. Each sample was assembled in two ways: the full arm assembles all reads, and each REFinder EC-N seeded arm assembles only the reads that REBEAN classifies as EC class N (N = 1–7), plus the reads recruited to their neighborhood by BBMap. For each EC class, four sets are defined: (I) full-arm proteins whose best Swiss-Prot hit carries an EC N number (DIAMOND); (II) the enzymes of I with at least one mapped read that REBEAN classifies as EC N (REBEAN filter); (III) EC-N seeded-arm proteins whose best hit carries an EC N number (DIAMOND); (IV) EC-N seeded-arm proteins with at least one mapped read that REBEAN assigns to EC N with a probability at or above the class-specific 90%-precision cutoff (REBEAN). Comparisons are class-matched: an EC-N seeded arm is compared only with the full arm’s EC-N enzymes, and an enzyme is shared (∩) only if both sets place it in the same class; \ denotes the enzymes in only one set. (a) Full versus seeded arm under the same homology annotation (II versus III), counted as distinct Swiss-Prot accessions (**Table 1, S1**) and summed over the seven classes, (b) Full arm with DIAMOND (I) versus REFinder seeded arm with REBEAN (IV), counted as enzyme clusters because REBEAN enzymes need not have a Swiss-Prot hit. Clusters are within-sample MMseqs2 clusters (90% identity, 80% coverage) pooling the full arm and all seven seeded arms, and each cluster is counted once per set. (c) Known and novel enzyme clusters in I, in IV, and in IV by EC class; a cluster is novel if no member has a Swiss-Prot hit at 30% identity or above. Boxes show the median and interquartile range (IQR), whiskers extend to the most extreme value within 1.5 × IQR, and points are individual samples; y axes in log scale.

### 3.2. REFinder identifies more enzymes than sequence similarity

Alignment-based methods can only annotate a protein when a similar sequence already exists in a reference database. Thus, they necessarily underestimate the enzymatic content of a metagenome. REFinder uses REBEAN for read annotation, i.e. prediction instead of homology. At our 90%-precision cutoff, which corresponds to approximately 30% recall in our benchmark ^21^, REFinder identified 1.2 fold more enzymes per oral sample and 2.1 fold more per murine sample than the full-arm using alignment, on average (**Tables 2, S2 & Figure 2b, S1b**). This meant that even for some of the best studied microbiomes, we could find over twenty thousand new, sequence-unique enzymes per sample. We expect that these numbers are higher in extreme (e.g. thermophilic) or hard-to-access (e.g. ocean floor) environments. One exception to this rule is the hydrolase (EC 3), where the full arm assembles more enzymes than the seeded arm. We hypothesize that hydrolases, the largest class in Swiss-Prot with diverse sequence families, favour homology annotation over REBEAN (set at 90% precision with low recall) that is trained for functional signatures ^21^. Note every assembled enzyme is a cluster representative of assembled genes at 90% sequence identity with 80% coverage (**Section 2.2.2**).

**Table 2:** Recovery of novel enzymes using REFinder seeded arm vs Full arm.

| EC | Class | No. of enzymes |  |  |  |  | No. of novel enzymes |  | Novel Overlap %<sup>†</sup> |
| --- | --- | --- | --- | --- | --- | --- | --- | --- | --- |
|  |  | Full arm | Seeded arm | Overlap | Unique to full arm | Unique to seeded arm | Full arm | Seeded arm |  |
| 1.* | Oxidoreductase | 9,178 ± 2,765 | 19,704 ± 6,777 | 4,165 ± 1,161 | 5,013 ± 1,634 | 15,540 ± 5,799 | 295 ± 104 | 5,941 ± 2,373 | 1.3 ± 0.2 |
| 2.* | Transferase | 26,758 ± 8,407 | 26,040 ± 8,665 | 10,600 ± 3,154 | 16,157 ± 5,365 | 15,439 ± 5,723 | 1,397 ± 487 | 5,784 ± 2,342 | 4.2 ± 0.6 |
| 3.* | Hydrolase | 22,261 ± 6,962 | 11,630 ± 3,992 | 4,307 ± 1,286 | 17,954 ± 5,750 | 7,323 ± 2,750 | 1,634 ± 569 | 2,571 ± 1,052 | 3.9 ± 0.9 |
| 4.* | Lyase | 5,370 ± 1,621 | 8,938 ± 3,187 | 2,125 ± 609 | 3,245 ± 1,021 | 6,813 ± 2,697 | 101 ± 33 | 2,415 ± 1,036 | 0.3 ± 0.1 |
| 5.* | Isomerase | 5,716 ± 1,840 | 11,171 ± 3,989 | 2,671 ± 785 | 3,044 ± 1,062 | 8,500 ± 3,370 | 156 ± 54 | 3,275 ± 1,379 | 0.6 ± 0.1 |
| 6.* | Ligase | 9,143 ± 3,031 | 10,543 ± 3,570 | 3,898 ± 1,155 | 5,245 ± 1,887 | 6,644 ± 2,627 | 210 ± 81 | 2,519 ± 1,050 | 1.2 ± 0.2 |
| 7.* | Translocase | 5,439 ± 1,687 | 14,325 ± 4,508 | 3,101 ± 885 | 2,337 ± 806 | 11,223 ± 3,792 | 152 ± 57 | 3,812 ± 1,430 | 0.7 ± 0.1 |

<sup>†</sup> The evaluation was carried out over 30 samples from PRJNA396840. Novel enzymes are the genes that code for proteins with less than 30% identity with Swiss-Prot database. An enzyme is a sequence representative of proteins (>90% identity), translated from each assembly that contain at least one REBEAN identified enzymatic read (seeded arm) or matches Swiss-Prot enzymes (full arm). Enzyme is considered novel if it shares less than 30% identity with proteins in Swiss-Prot database. Two enzymes from two different assemblies are said to overlap if they cluster together (≥ 90% identity with ≥ 80% of both sequences covered).

Novelty of an identified enzyme is defined by lack of homology to known protein space (**Section 2.3.2**). In this study, to be considered extant (already annotated) an enzyme needed a Swiss-Prot hit; a sequence with no hit or only with hits below 30% sequence identity counted as novel. A substantial number of REFinder-assembled enzymes are sequence-novel, ranging from 22.1 to 30.1% and 36.4 to 45.0% in oral saliva and murine gut, respectively (**Figure 2c, S1c & Tables 2, S2)**. In comparison, novel enzymes are a minority of what similarity searches can recover, i.e. 1.9% to 7.3% of full-assembly enzymes per EC class in saliva and 2.5% to 9.6% in the murine gut.

Using REBEAN instead of DIAMOND alignment ^36^ in the full arm resulted in a much higher novel enzyme fraction: 22.1% to 29.9% per EC class in saliva and 36.6% to 44.6% in the murine gut – similar in fraction to REFinder’s seeded arm (**Table S3**), highlighting the usefulness of REBEAN, the core of REFinder pipeline, in discovery of novel enzymes.

As described above, the comparison between assembly arms was based on the number of enzyme clusters defined at 90% sequence identity, a cut-off that prevents duplicated and fragmented sequences from inflating the count. A complementary view is obtained by clustering at 30% identity, which groups related sequences together so that each distinct enzyme group is counted once. We therefore repeated the comparison at this threshold. Under this coarser definition, the seeded arm recovers 1.5-fold (saliva) and 3.1-fold (murine gut) more enzyme clusters per sample than the full arm. The difference widens because relaxing the cut-off from 90% to 30% reduces the seeded-arm cluster count by 28.5 ± 5.8% and 38.4 ± 7.2% in saliva and murine gut, respectively, whereas the corresponding reductions in the full arm are larger, at 43.2 ± 5.4% and 58.6 ± 5.6% (**Tables 2 and S2-S3; Figures 2b and S1b-S3b**). The fraction of novel enzymes in the seeded arm also increased, from 26.7% to 34.5% in saliva and from 42.9% to 51.7% in murine gut. The findings reported above are therefore robust to the definition used for enzyme distinctiveness.

### 3.3. REFinder novel proteins have high structural similarity to known enzymes

Because REFinder-only calls cannot be confirmed by sequence alignment, we asked whether they correspond to real catalytic proteins by way of structure alignment. We screened the 23,190 novel EC1 enzymes, each representative of a corresponding protein cluster and each with the best Swiss-Prot hit below 30% identity, from two saliva samples (SRR5892215 & SRR5892216) for reliable ESM2 650M ^37^ embeddings (RNS ^38^. RNS is a measure of embedding uncertainty that affects downstream tasks including structure prediction, and our earlier work has shown consistent blind spots among pLM variants. We predicted structures (ESMFold average pLDDT 84.4 ± 8.9) for the 13,745 enzymes (with RNS = 0). Of these 5,180 (≥0.7 TM-score) and 772 (≥0.9 TM-score) proteins structurally aligned with EC-annotated PDB chains ^18, 44, 45^ despite negligible sequence identity i.e. 37.7% of proteins have considerable structural matches without sequence homology.

In addition to protein structure prediction, REFinder runs searches against Pfam HMMs that mapped 10,664 (77.5%) of the 13,745 to known (69.8%) and unknown (7.7%) domain families. Out of the 5,180 recovered enzymes that share structural similarity with PDB enzymes, 87.9% map to 1,341 distinct known and 1.6% to 101 distinct unknown Pfam domains (DUFs). The remaining 10.5% (545 of 5,180) of novel enzymes were not recoverable through hmm searches. Further illustrating the usefulness of REFinder’s activity-driven targeted assembly for exploration of microbiomes.

Figure 3. shows four such proteins matched and aligned against experimentally studied PDB structures ^18, 44, 46^ with known oxidoreductase EC annotation ^45^ and no detectable matches in Swiss-Prot and Pfam-A. NODE_21166 (168 aa) matched spinach ferredoxin–nitrite reductase (PDB 2AKJ_A; TM-score 0.753, 5.9% sequence identity; EC 1.7.7.1). NODE_39435 (308 aa) matched the dimethylglycine oxidase of Arthrobacter globiformis (1PJ6_A; TM-score 0.779, 7.2% identity; EC 1.5.3.10). NODE_56285 (180 aa) matched the F(420)-dependent glucose-6-phosphate dehydrogenase (FGD1) of Mycobacterium tuberculosis (3C8N_D; TM-score 0.758, 11.6% identity; EC 1.1.98.2). NODE_31123 (378 aa) matched the γ-carotene desaturase of Nonlabens dokdonensis (4REP_A; TM-score 0.833, 12.7% identity; EC 1.3.99.27).

At 6-13% sequence identity with the corresponding PDB sequences, these matches fall far below the range in which homology transfer is reliable, so these proteins would be deemed homology-invisible, while adopting folds of well-characterized enzymes. Note that the structural correspondence establishes fold rather than activity and does not by itself demonstrate catalysis, but it shows that a substantial share of the homology-invisible set of proteins is neither a result of assembly nor an artifact of enzyme classification. The most prominent value of REFinder is that it retains these proteins at a fraction of the assembly cost, not that it proves these proteins are enzymes.

**Figure 3.**
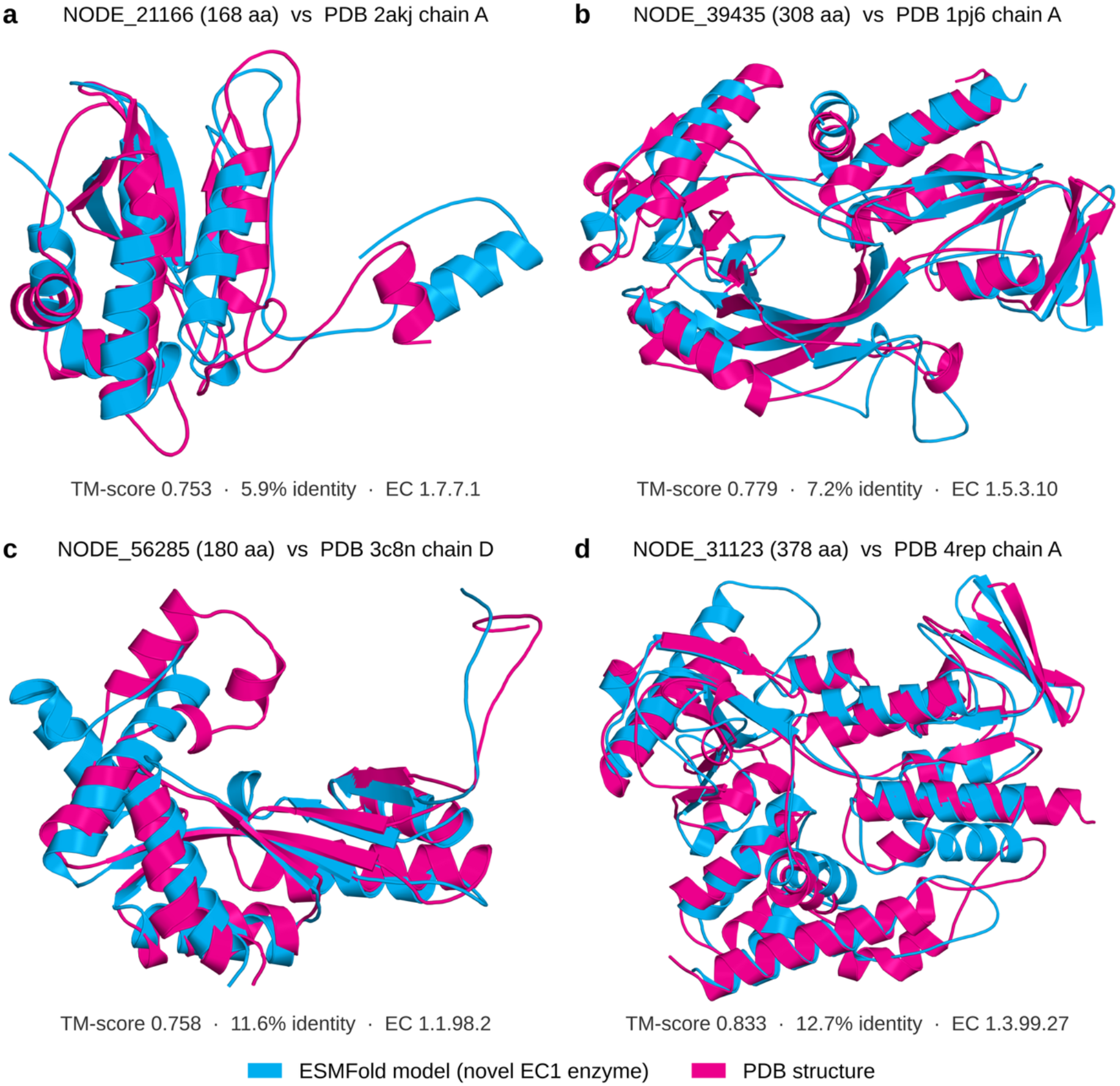
REFinder identified novel, homology-invisible enzymes show structural similarity to characterized oxidoreductases. ESMFold models (cyan) of four novel enzymes recovered by the seeded assembly of saliva metagenomes, superposed on their best EC-annotated PDB chains (magenta). Each protein was assigned to EC1 by REBEAN, has a reliable embedding (RNS = 0), has no Swiss-Prot hit at 30% identity or above, and has no Pfam-A match (neither at the gathering thresholds nor at E ≤ 10^−5^); (a) NODE_21166 (168 aa; SRR5892215) against spinach ferredoxin–nitrite reductase (PDB 2akj chain A, 535 aa): TM-score 0.753, 5.9% sequence identity, RMSD 4.43 Å over 152 residues, EC 1.7.7.1. (b) NODE_39435 (308 aa; SRR5892216) against Arthrobacter globiformis N,N-dimethylglycine oxidase (PDB 1pj6 chain A, 828 aa): TM-score 0.779, 7.2% identity, RMSD 4.49 Å over 272 residues, EC 1.5.3.10. (c) NODE_56285 (180 aa; SRR5892216) against Mycobacterium tuberculosis F420-dependent glucose-6-phosphate dehydrogenase FGD1 (PDB 3c8n chain D, 330 aa): TM-score 0.758, 11.6% identity, RMSD 3.84 Å over 160 residues, EC 1.1.98.2. (d) NODE_31123 (378 aa; SRR5892216) against Nonlabens dokdonensis γ-carotene desaturase (PDB 4rep chain A, 489 aa): TM-score 0.833, 12.7% identity, RMSD 3.40 Å over 344 residues, EC 1.3.99.27. Models are confident (mean pLDDT 87–92). TM-scores and sequence identities are from Foldseek (TM-align mode) over the structural alignment.

### 3.4. Assembling genes at low read-depth

A class-focused assembly should be able to reconstruct genes so sparsely covered by sequencing that they are unlikely to emerge from the whole community where they compete against the entire read pool. To test this hypothesis, we counted the trimmed sample reads that mapped to each assembled gene and compared the coverage of arm-unique enzymes. Seed-only enzymes were, on average, of lower read depth than assembly-only enzymes in all seven EC classes across both datasets; median coverage was 17-27 reads per enzyme for seed-only vs. 24-51 for assembly-only in the saliva set, and 22-49 vs. 30-95, respectively, in the murine gut, a median ratio of seed-to-full assembly proteins of 0.52-0.73 (**Figure S4, S5**). Seeding therefore uniquely recovers genes that are drawn from the sparsely covered assembly tail, i.e. the regime in which enzyme mining is hardest. We hypothesize that restricting assembly to class-routed reads may simplify the de Bruijn graph used in assembly by removing unrelated sequence and improving contiguity for the targeted class. This may also stem from low depth reads assembled into shorter contigs and incomplete genes (**Figure S5**). Though not established herein, these hypotheses are testable by controlled down-sampling and by direct measurement of graph complexity. In addition, we observed seeded assemblies were enriched with enzymatic reads. In the seeded assemblies, enzyme genes accounted for a median of 9.2% (saliva) and 5.4% (murine gut) of the mapped reads when annotated by DIAMOND, and 33.1% and 41.1% when annotated by REBEAN. In the full assembly, the corresponding shares were 5.2% and 4.0% for DIAMOND-annotated enzyme genes (**Figure S5**) and 19.7% and 32.8% for REBEAN-annotated ones.

### 3.5. Computational cost

Assembly compute cost dominated both study arms. Per saliva sample, metaSPAdes accounted for 10.3 ± 5.6 of the 10.9 ± 5.7 CPU-hours of a full assembly; in the murine gut it accounted for 64.1 ± 13.9 of 65.5 ± 14.0 CPU-hours (**Table S5**). Because each seed arm assembled only its class-routed reads, their costs fell below the full arm in most classes: the saliva EC-*N* seed arms ranged from 1.7 ± 0.9 CPU-hours for EC7 (0.16 times the full arm) to 8.0 ± 4.6 CPU-hours for EC2 (0.73 times the full arm), for a maximum saving of 6.4-fold. The murine seed arm ranged from 18.5 ± 6.1 CPU-hours for EC7 (0.28 times full assembly) to 71.3 ± 16.7 CPU-hours for EC2 (1.09 times full assembly). Note that murine EC2, the class to which REBEAN routed the most reads, was the sole exception to cost savings – every other EC class in every sample was cheaper in seeded than in full assembly states. Consistent with prior targeted assemblers ^25^, restricting assembly to a single routed class reduces cost relative to whole-community assembly. Note that recovering the complete enzyme catalogue is a different question than running seven separate seed arms, one per EC class. A single assembly of all enzymatic reads is expected to cost less than the sum across ECs or the full assembly, but this runtime was not evaluated here as it is not the scope for a seeded assembly framework. In addition, this accounting excluded the GPU usage for REBEAN enzyme annotation of saliva (6.5±3.2M reads per sample) and gut (70.1±13.3M reads) samples (0.13±0.06 h and 1.4±0.3 per sample, respectively; **Table S5)**.

## 4. Conclusions

This is, to our knowledge, the first implementation of an alignment-free, language model-guided, targeted, enzyme discovery pipeline. REFinder reconstructs known enzymes comparably to full assembly while recovering large numbers of sequence-novel enzymes that homology cannot reach, at lower assembly cost per targeted class.

The present system routes metagenomic reads at the top level of the EC hierarchy, which is coarse: a class such as EC7, which includes enzymes that transport specific substrates, is still an aggregate of many distinct molecular activities. Routing on finer EC sub-classes would further concentrate assembly, reducing cost and sharpening recovery of a targeted activity. While it is the natural next step for this work, we note that discovery of novel (non-EC-elucidated) enzymatic activities may be a direct product of coarse-grained classification presented here.

Three limitations bound these conclusions. First, here we evaluated two well-studied communities/datasets, so generalization to high-diversity soil and marine environments remains to be shown. Second, cost savings are class dependent. In fact, we found one outlier: for murine EC2, the class recruiting the most sample reads, seeding cost slightly more than full assembly. Most importantly, our compute accounting excludes the cost of running REBEAN over the read sets. We aim to optimize its speed in later versions of the pipeline.

## Supporting information

SOM

## 5. Acknowledgments

## 6. Author contributions

Conceptualization: RP, YB; Development & Methodology: RP; Visualization: RP, YB; Supervision: YB; Writing and revisions: RP, YB;

## 8. Data and materials availability

The pipeline is available at https://bitbucket.org/bromberglab/NF-REFinder. The underlying standalone REBEAN package can be found at https://bitbucket.org/bromberglab/rebeanpkg.

