## Supplementary material for "REFinder: Mining new enzymes from metagenomes": SOM

R. Prabakaran[1] and Yana Bromberg[1,2†],

[1]*Department of Biology, Emory University,*
*Atlanta, GA 30322, USA*

[2]*Department of Computer Science, Emory University,*
*Atlanta, GA 30322, USA*

**

#### Supplementary Tables

##### Table S1: Recovery of known enzymes from the murine gut metagenomes

| EC | Class | No. of distinct Swiss-Prot Enzymes identified | | | | | Overlap % |
|---|---|---|---|---|---|---|---|
| | | Full arm | Full arm with REBEAN filter† | Seeded arm | Unique to Full arm | Unique to Seeded arm | |
| 1.* | Oxidoreductase | 3,221±448 | 2,870±410 | 2,865±407 | 222±45 | 218±40 | 85.8±0.9 |
| 2.* | Transferase | 9,335±1472 | 8,516±1362 | 8,507±1354 | 597±124 | 588±116 | 87.1±0.7 |
| 3.* | Hydrolase | 6,812±998 | 5,182±778 | 5,265±783 | 599±101 | 682±109 | 78.2±1.0 |
| 4.* | Lyase | 2,119±350 | 1,679±270 | 1,716±276 | 198±39 | 234±47 | 77.5±1.6 |
| 5.* | Isomerase | 1,829±315 | 1,492±256 | 1,532±254 | 161±35 | 201±36 | 78.6±1.2 |
| 6.* | Ligase | 2,884±656 | 2,257±512 | 2,404±535 | 282±68 | 429±95 | 73.5±1.1 |
| 7.* | Translocase | 1,697±271 | 1,486±249 | 1,535±253 | 140±24 | 189±32 | 80.3±1.1 |

† To ensure a standardized efficiency comparison between the Seeded and Full arm assemblies, we restricted the analysis to genes containing at least one REBEAN-predicted enzymatic read. The mean and standard deviation over the 20 samples (PRJNA504846) is reported.

#### Table S2: Recovery of novel enzymes using seeded arm vs Full arm

| EC | No. of enzymes | | | | | No. of novel enzymes | | Novel overlap %† |
|---|---|---|---|---|---|---|---|---|
| | Full arm | Seeded arm | Overlap | Unique to Full arm | Unique to Seeded arm | Full arm | Seeded arm | |
| Oral saliva (n = 30) | | | | | | | | |
| 1.* | 9,178 ± 2,765 | 19,704 ± 6,777 | 4,165 ± 1,161 | 5,013 ± 1,634 | 15,540 ± 5,799 | 295 ± 104 | 5,941 ± 2,373 | 1.3 ± 0.2 |
| 2.* | 26,758 ± 8,407 | 26,040 ± 8,665 | 10,600 ± 3,154 | 16,157 ± 5,365 | 15,439 ± 5,723 | 1,397 ± 487 | 5,784 ± 2,342 | 4.2 ± 0.6 |
| 3.* | 22,261 ± 6,962 | 11,630 ± 3,992 | 4,307 ± 1,286 | 17,954 ± 5,750 | 7,323 ± 2,750 | 1,634 ± 569 | 2,571 ± 1,052 | 3.9 ± 0.9 |
| 4.* | 5,370 ± 1,621 | 8,938 ± 3,187 | 2,125 ± 609 | 3,245 ± 1,021 | 6,813 ± 2,697 | 101 ± 33 | 2,415 ± 1,036 | 0.3 ± 0.1 |
| 5.* | 5,716 ± 1,840 | 11,171 ± 3,989 | 2,671 ± 785 | 3,044 ± 1,062 | 8,500 ± 3,370 | 156 ± 54 | 3,275 ± 1,379 | 0.6 ± 0.1 |
| 6.* | 9,143 ± 3,031 | 10,543 ± 3,570 | 3,898 ± 1,155 | 5,245 ± 1,887 | 6,644 ± 2,627 | 210 ± 81 | 2,519 ± 1,050 | 1.2 ± 0.2 |
| 7.* | 5,439 ± 1,687 | 14,325 ± 4,508 | 3,101 ± 885 | 2,337 ± 806 | 11,223 ± 3,792 | 152 ± 57 | 3,812 ± 1,430 | 0.7 ± 0.1 |
| Murine gut (n = 20) | | | | | | | | |
| 1.* | 19,181 ± 5,781 | 75,265 ± 16,407 | 10,370 ± 2,752 | 8,811 ± 3,077 | 64,895 ± 13,859 | 1,120 ± 281 | 33,867 ± 7,158 | 1.7 ± 0.2 |
| 2.* | 58,734 ± 18,134 | 93,222 ± 21,583 | 28,532 ± 7,924 | 30,202 ± 10,334 | 64,690 ± 14,087 | 4,319 ± 1,163 | 34,338 ± 7,257 | 4.8 ± 0.3 |
| 3.* | 46,208 ± 14,260 | 39,747 ± 8,903 | 11,187 ± 2,719 | 35,021 ± 11,652 | 28,560 ± 6,244 | 4,433 ± 1,164 | 14,467 ± 2,981 | 5.6 ± 0.4 |
| 4.* | 10,171 ± 3,045 | 37,249 ± 7,845 | 4,421 ± 1,178 | 5,750 ± 1,877 | 32,828 ± 6,818 | 440 ± 120 | 16,304 ± 3,340 | 0.4 ± 0.1 |
| 5.* | 11,090 ± 3,594 | 41,070 ± 8,855 | 5,729 ± 1,685 | 5,361 ± 1,919 | 35,341 ± 7,418 | 385 ± 104 | 18,449 ± 3,851 | 0.4 ± 0.1 |
| 6.* | 15,618 ± 5,397 | 35,938 ± 7,922 | 7,461 ± 2,312 | 8,157 ± 3,094 | 28,477 ± 5,948 | 396 ± 124 | 14,907 ± 3,026 | 0.6 ± 0.1 |
| 7.* | 10,455 ± 3,289 | 45,142 ± 10,460 | 6,448 ± 1,909 | 4,007 ± 1,386 | 38,694 ± 8,701 | 263 ± 87 | 17,695 ± 3,861 | 0.4 ± 0.1 |

† An enzyme is a sequence representative of proteins (>90% identity), translated from each assembly that contain at least one REBEAN identified enzymatic read (seeded arm) or matches Swiss-Prot enzymes (full arm). Enzyme is considered novel if it shares less than 30% identity with proteins in Swiss-Prot database. Two enzymes from two different assemblies are said to overlap if they cluster together (≥ 90% identity with ≥ 80% of both sequences covered). Mean and standard deviation across samples are reported.

#### Table S3: REBEAN identifies novel enzymes in both assembly pipelines

| EC | Class | No. of REBEAN identified enzymes at 90% precision | | | | | No. of novel enzymes | | Overlap %† |
|---|---|---|---|---|---|---|---|---|---|
| | | Full arm (REBEAN) | Seeded arm | Overlap | Unique to Full arm (REBEAN) | Unique to Seeded arm | Full arm (REBEAN) | Seeded arm | |
| Oral saliva (n = 30) | | | | | | | | | |
| 1.* | Oxidoreductase | 19,866±6815 | 19,704±6777 | 14,568±4854 | 5,298±2025 | 5,136±1981 | 5,932±2372 | 5,941±2373 | 58.3 |
| 2.* | Transferase | 26,214±8718 | 26,040±8665 | 21,418±6882 | 4,797±1893 | 4,622±1836 | 5,792±2340 | 5,784±2342 | 69.5 |
| 3.* | Hydrolase | 11,726±4007 | 11,630±3992 | 7,910±2557 | 3,816±1472 | 3,721±1457 | 2,591±1056 | 2,571±1052 | 51.2 |
| 4.* | Lyase | 9,024±3221 | 8,938±3187 | 4,992±1610 | 4,031±1640 | 3,946±1610 | 2,371±1021 | 2,415±1036 | 38.5 |
| 5.* | Isomerase | 11,272±4018 | 11,171±3989 | 6,377±2064 | 4,895±1984 | 4,794±1959 | 3,220±1355 | 3,275±1379 | 39.7 |
| 6.* | Ligase | 10,610±3591 | 10,543±3570 | 6,409±1971 | 4,200±1671 | 4,133±1658 | 2,476±1036 | 2,519±1050 | 43.5 |
| 7.* | Translocase | 14,377±4496 | 14,325±4508 | 8,645±2417 | 5,732±2145 | 5,680±2174 | 3,742±1407 | 3,812±1430 | 43.1 |
| Murine gut (n = 20) | | | | | | | | | |
| 1.* | Oxidoreductase | 76,993±16946 | 75,265±16407 | 58,723±11820 | 18,270±5782 | 16,542±5161 | 34,332±7309 | 33,867±7158 | 62.8 |
| 2.* | Transferase | 94,818±22133 | 93,222±21583 | 79,093±17345 | 15,725±5150 | 14,129±4557 | 34,902±7428 | 34,338±7257 | 72.6 |
| 3.* | Hydrolase | 40,751±9255 | 39,747±8903 | 28,734±5899 | 12,017±3696 | 11,013±3306 | 14,821±3091 | 14,467±2981 | 55.5 |
| 4.* | Lyase | 38,302±8167 | 37,249±7845 | 20,999±4122 | 17,303±4495 | 16,250±4106 | 16,282±3376 | 16,304±3340 | 38.5 |
| 5.* | Isomerase | 42,301±9264 | 41,070±8855 | 24,010±4757 | 18,291±4936 | 17,060±4467 | 18,413±3897 | 18,449±3851 | 40.4 |
| 6.* | Ligase | 36,915±8267 | 35,938±7922 | 20,834±4195 | 16,080±4294 | 15,103±3920 | 14,960±3056 | 14,907±3026 | 40.1 |
| 7.* | Translocase | 46,436±10923 | 45,142±10460 | 25,122±5332 | 21,314±5695 | 20,020±5207 | 17,677±3872 | 17,695±3861 | 37.8 |

† An enzyme is a sequence representative of proteins (>90% identity), translated from each assembly that contain at least one REBEAN identified enzymatic read. Enzyme is considered novel if it shares less than 30% identity with proteins in Swiss-Prot database. Two enzymes from two different assemblies are said to overlap if they cluster together (≥ 90% identity with ≥ 80% of both sequences covered). Mean and standard deviation across samples are reported.

#### Table S4: Recovery of novel enzymes clusters (> 30%) using seeded arm vs Full arm

| EC | No. of enzymes | | | | | No. of novel enzymes | | Overlap %[†] |
|---|---|---|---|---|---|---|---|---|
| | Full arm | Seeded arm | Overlap | Unique to Full arm | Unique to Seeded arm | Full arm | Seeded arm | |
| Oral saliva (n = 30) | | | | | | | | |
| 1.* | 5,411 ± 1,294 | 14,276 ± 4,226 | 2,404 ± 516 | 3,007 ± 787 | 11,872 ± 3,820 | 172 ± 49 | 5,015 ± 1,794 | 0.9 ± 0.1 |
| 2.* | 14,701 ± 3,696 | 16,330 ± 4,541 | 5,367 ± 1,195 | 9,334 ± 2,522 | 10,962 ± 3,470 | 779 ± 201 | 4,768 ± 1,710 | 3.2 ± 0.4 |
| 3.* | 13,631 ± 3,571 | 8,053 ± 2,444 | 2,699 ± 668 | 10,931 ± 2,931 | 5,353 ± 1,831 | 971 ± 269 | 2,197 ± 824 | 3.3 ± 0.7 |
| 4.* | 3,039 ± 713 | 6,929 ± 2,313 | 1,178 ± 251 | 1,861 ± 466 | 5,751 ± 2,140 | 63 ± 17 | 2,164 ± 877 | 0.2 ± 0.1 |
| 5.* | 2,898 ± 755 | 8,422 ± 2,840 | 1,268 ± 279 | 1,630 ± 480 | 7,154 ± 2,639 | 92 ± 28 | 2,908 ± 1,152 | 0.4 ± 0.1 |
| 6.* | 4,995 ± 1,361 | 7,555 ± 2,383 | 2,130 ± 483 | 2,865 ± 894 | 5,425 ± 2,026 | 118 ± 38 | 2,245 ± 879 | 0.8 ± 0.2 |
| 7.* | 2,854 ± 711 | 9,409 ± 2,766 | 1,605 ± 367 | 1,249 ± 350 | 7,805 ± 2,487 | 90 ± 26 | 3,290 ± 1,129 | 0.5 ± 0.1 |
| Murine gut (n = 20) | | | | | | | | |
| 1.* | 8,238 ± 1,946 | 43,084 ± 8,985 | 4,240 ± 978 | 3,997 ± 987 | 38,844 ± 8,059 | 351 ± 81 | 22,237 ± 4,524 | 0.8 ± 0.1 |
| 2.* | 23,079 ± 5,338 | 46,170 ± 9,506 | 10,261 ± 2,322 | 12,818 ± 3,080 | 35,909 ± 7,300 | 1,426 ± 312 | 21,476 ± 4,284 | 2.5 ± 0.2 |
| 3.* | 21,367 ± 5,469 | 23,349 ± 4,693 | 5,268 ± 1,262 | 16,099 ± 4,234 | 18,081 ± 3,497 | 1,668 ± 376 | 9,984 ± 1,934 | 3.6 ± 0.4 |
| 4.* | 4,157 ± 946 | 25,717 ± 5,007 | 1,780 ± 374 | 2,377 ± 575 | 23,937 ± 4,674 | 201 ± 46 | 12,297 ± 2,400 | 0.3 ± 0.1 |
| 5.* | 4,063 ± 990 | 27,875 ± 5,526 | 2,022 ± 463 | 2,040 ± 532 | 25,853 ± 5,121 | 142 ± 33 | 14,008 ± 2,792 | 0.2 ± 0.0 |
| 6.* | 5,750 ± 1,449 | 23,934 ± 4,658 | 3,039 ± 758 | 2,710 ± 696 | 20,894 ± 4,027 | 123 ± 36 | 11,319 ± 2,194 | 0.3 ± 0.1 |
| 7.* | 3,988 ± 996 | 26,675 ± 5,447 | 2,540 ± 650 | 1,448 ± 352 | 24,134 ± 4,865 | 118 ± 35 | 13,046 ± 2,681 | 0.2 ± 0.0 |

[†] An enzyme cluster is a sequence representative of proteins (>30% identity), translated from each assembly that contain at least one REBEAN identified enzymatic read (seeded arm) or matches Swiss-Prot enzymes (full arm). Enzyme is considered novel if it shares less than 30% identity with proteins in Swiss-Prot database. Two enzymes from two different assemblies are said to overlap if they cluster together (≥ 30% identity with ≥ 80% of both sequences covered). Mean and standard deviation across samples are reported.

#### Table S5: Computational cost per sample by pipeline stage

| arm | Dataset | Time taken (CPU-h) | | | | |
|---|---|---|---|---|---|---|
| | | Bait (BBMap) | Assembly (MetaSPAdes) | Gene calling (Prodigal) | Alignment to Swiss-Prot (DIAMOND) | Total |
| Full | Saliva | 0.0±0.0 | 10.3±5.6 | 0.1±0.0 | 0.5±0.1 | 10.9±5.7 |
| Seeded EC 1* | Saliva | 0.9±0.6 | 5.3±3.4 | 0.0±0.0 | 0.3±0.1 | 6.6±4.0 |
| Seeded EC 2.* | Saliva | 1.1±0.7 | 6.5±3.9 | 0.1±0.0 | 0.4±0.1 | 8.0±4.6 |
| Seeded EC 3.* | Saliva | 0.8±0.5 | 3.5±2.2 | 0.0±0.0 | 0.3±0.1 | 4.6±2.7 |
| Seeded EC 4.* | Saliva | 0.5±0.3 | 1.6±1.1 | 0.0±0.0 | 0.2±0.0 | 2.4±1.4 |
| Seeded EC 5.* | Saliva | 0.6±0.4 | 2.3±1.5 | 0.0±0.0 | 0.2±0.0 | 3.1±1.9 |
| Seeded EC 6.* | Saliva | 0.5±0.4 | 1.9±1.1 | 0.0±0.0 | 0.2±0.0 | 2.7±1.5 |
| Seeded EC 7.* | Saliva | 0.4±0.2 | 1.1±0.7 | 0.0±0.0 | 0.2±0.0 | 1.7±0.9 |
| Full | Murine | 0.0±0.0 | 64.1±13.9 | 0.5±0.1 | 0.9±0.2 | 65.5±14.0 |
| Seeded EC 1* | Murine | 11.7±3.9 | 42.9±8.2 | 0.3±0.1 | 0.6±0.1 | 55.4±11.4 |
| Seeded EC 2.* | Murine | 17.4±6.1 | 52.8±11.2 | 0.4±0.1 | 0.7±0.1 | 71.3±16.7 |
| Seeded EC 3.* | Murine | 8.8±2.9 | 35.9±8.4 | 0.2±0.0 | 0.5±0.1 | 45.4±10.9 |
| Seeded EC 4.* | Murine | 5.2±1.5 | 25.4±7.4 | 0.1±0.0 | 0.3±0.0 | 31.1±8.8 |
| Seeded EC 5.* | Murine | 5.7±1.9 | 23.6±6.8 | 0.1±0.0 | 0.4±0.1 | 29.8±8.5 |
| Seeded EC 6.* | Murine | 5.3±1.8 | 21.1±5.8 | 0.1±0.0 | 0.4±0.1 | 26.9±7.5 |
| Seeded EC 7.* | Murine | 4.1±1.3 | 14.1±4.8 | 0.1±0.0 | 0.3±0.0 | 18.5±6.1 |

† REBEAN EC predictions for each sample required 0.13±0.06 h and 1.4±0.3 h for saliva (6.5±3.2M reads per sample) and gut (70.1±13.3M reads) samples, respectively on a NVIDIA RTX 6000 Ada 48GB card using 8 CPU cores. Mean and standard deviation across samples are reported.

### Supplementary Figures

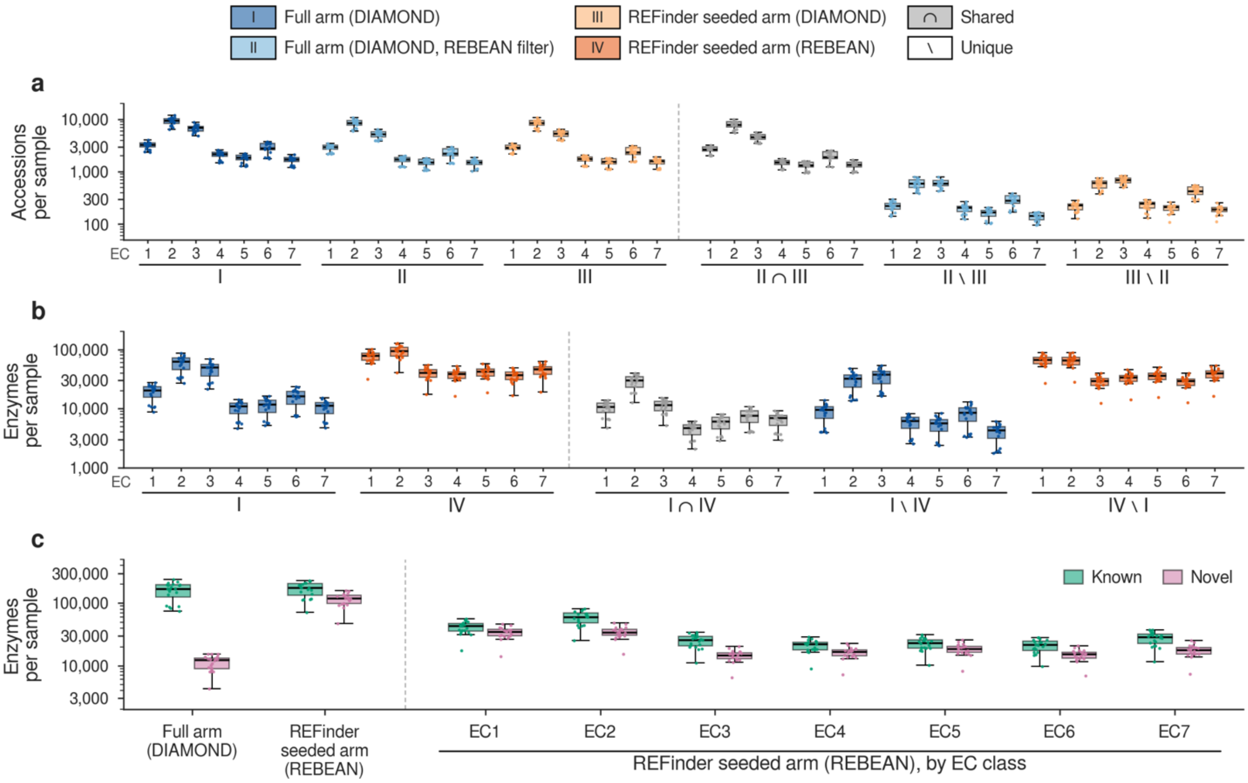


**Figure S1**: **Enzymes recovered from murine gut metagenomes by full arm and REFinder's seeded arm.** Each sample was assembled in two ways: the full arm assembles all reads, and each REFinder EC-N seeded arm assembles only the reads that REBEAN classifies as EC class N (N = 1–7), plus the reads recruited to their neighborhood by BBMap. For each EC class, four sets are defined: (I) full-arm proteins whose best Swiss-Prot hit carries an EC N number (DIAMOND); (II) the enzymes of I with at least one mapped read that REBEAN classifies as EC N (REBEAN filter); (III) EC-N seeded-arm proteins whose best hit carries an EC N number (DIAMOND); (IV) EC-N seeded-arm proteins with at least one mapped read that REBEAN assigns to EC N with a probability at or above the class-specific 90%-precision cutoff (REBEAN). Comparisons are class-matched: an EC-N seeded arm is compared only with the full arm's EC-N enzymes, and an enzyme is shared (∩) only if both sets place it in the same class; \ denotes the enzymes in only one set. (a) Full versus seeded arm under the same homology annotation (II versus III), counted as distinct Swiss-Prot accessions (**Table 1, S1**) and summed over the seven classes, (b) Full arm with DIAMOND (I) versus REFinder seeded arm with REBEAN (IV), counted as enzyme clusters because REBEAN enzymes need not have a Swiss-Prot hit. Clusters are within-sample MMseqs2 clusters (90% identity, 80% coverage) pooling the full arm and all seven seeded arms, and each cluster is counted once per set. (c) Known and novel enzyme clusters in I, in IV, and in IV by EC class; a cluster is novel if no member has a Swiss-Prot hit at 30% identity or above. Boxes show the median and interquartile range (IQR), whiskers extend to the most extreme value within 1.5 × IQR, and points are individual samples; y axes in log scale.

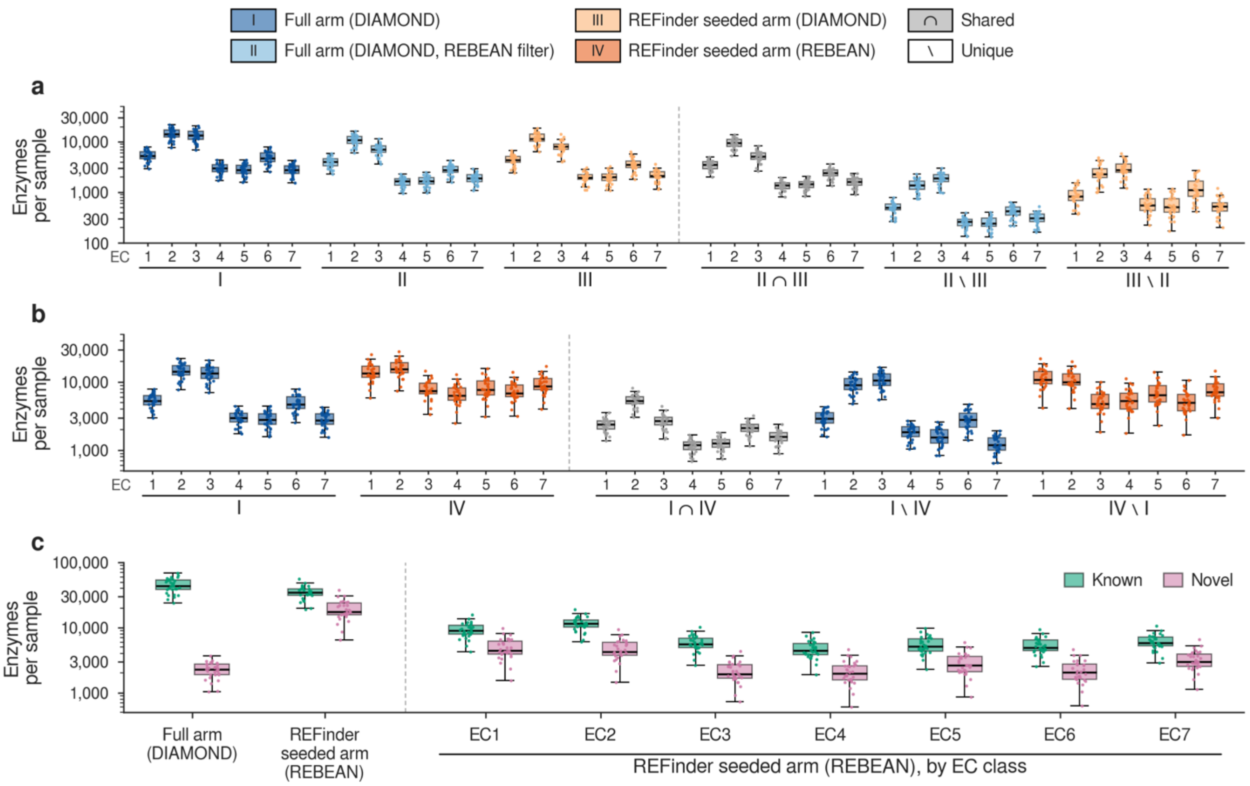


**Figure S2**: **Enzyme clusters recovered from oral saliva metagenomes by full arm and REFinder's seeded arm.** Each sample was assembled in two ways: the full arm assembles all reads, and each REFinder EC-N seeded arm assembles only the reads that REBEAN classifies as EC class N (N = 1–7), plus the reads recruited to their neighborhood by BBMap. For each EC class, four sets are defined: (I) full-arm proteins whose best Swiss-Prot hit carries an EC N number (DIAMOND); (II) the enzymes of I with at least one mapped read that REBEAN classifies as EC N (REBEAN filter); (III) EC-N seeded-arm proteins whose best hit carries an EC N number (DIAMOND); (IV) EC-N seeded-arm proteins with at least one mapped read that REBEAN assigns to EC N with a probability at or above the class-specific 90%-precision cutoff (REBEAN). Comparisons are class-matched: an EC-N seeded arm is compared only with the full arm's EC-N enzymes, and an enzyme is shared (∩) only if both sets place it in the same class; \ denotes the enzymes in only one set.

All counts reflect clusters within-sample MMseqs2 clusters (30% identity, 80% coverage) pooling the full arm and all seven seeded arms, and each cluster is counted once per set. (a) Full versus seeded arm under the same homology annotation (II versus III), counted as distinct Swiss-Prot accessions (**Table 1, S1**) and summed over the seven classes, (b) Full arm with DIAMOND (I) versus REFinder seeded arm with REBEAN (IV), counted as enzyme clusters because REBEAN enzymes need not have a Swiss-Prot hit. (c) Known and novel enzyme clusters in I, in IV, and in IV by EC class; a cluster is novel if no member has a Swiss-Prot hit at 30% identity or above. Boxes show the median and interquartile range (IQR), whiskers extend to the most extreme value within 1.5 × IQR, and points are individual samples; y axes in log scale.

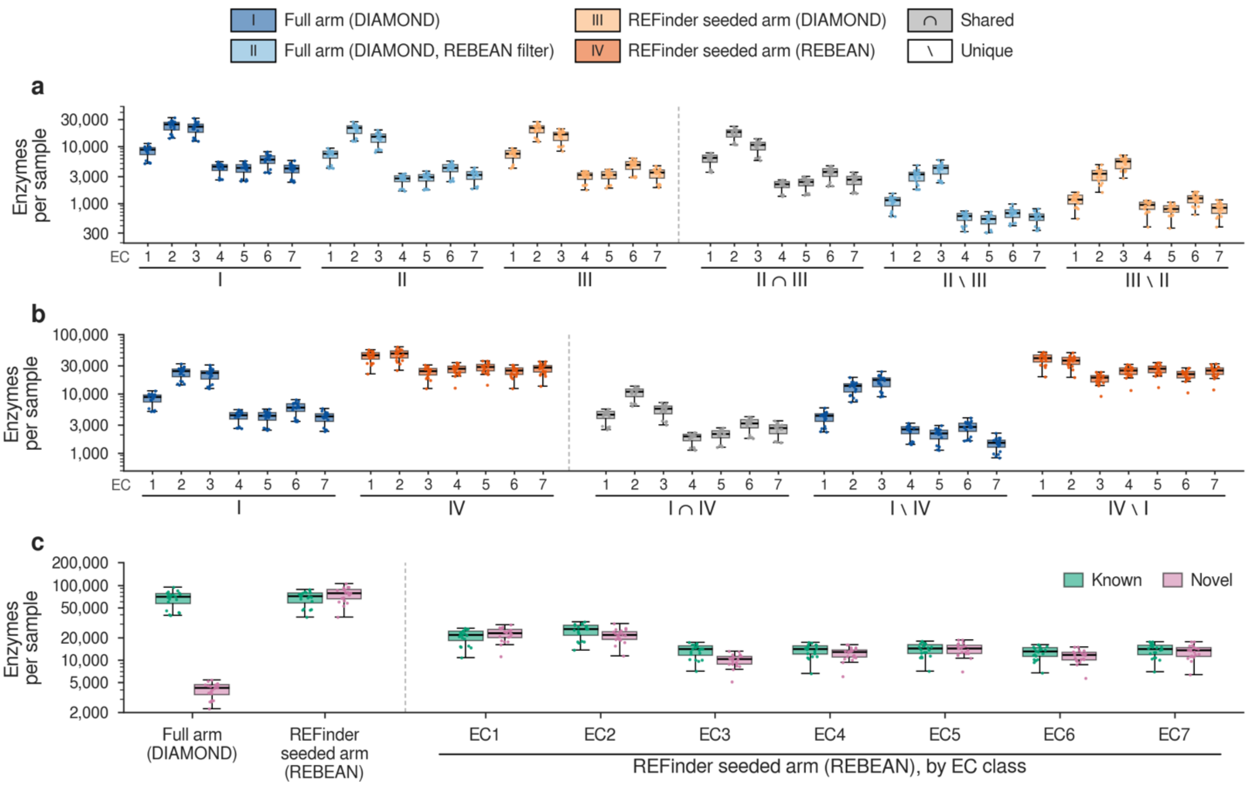


**Figure S3**: **Enzyme clusters recovered from murine gut metagenomes by full arm and REFinder's seeded arm.** Each sample was assembled in two ways: the full arm assembles all reads, and each REFinder EC-N seeded arm assembles only the reads that REBEAN classifies as EC class N (N = 1–7), plus the reads recruited to their neighborhood by BBMap. For each EC class, four sets are defined: (I) full-arm proteins whose best Swiss-Prot hit carries an EC N number (DIAMOND); (II) the enzymes of I with at least one mapped read that REBEAN classifies as EC N (REBEAN filter); (III) EC-N seeded-arm proteins whose best hit carries an EC N number (DIAMOND); (IV) EC-N seeded-arm proteins with at least one mapped read that REBEAN assigns to EC N with a probability at or above the class-specific 90%-precision cutoff (REBEAN). Comparisons are class-matched: an EC-N seeded arm is compared only with the full arm's EC-N enzymes, and an enzyme is shared (∩) only if both sets place it in the same class; \ denotes the enzymes in only one set.

All counts reflect clusters within-sample MMseqs2 clusters (30% identity, 80% coverage) pooling the full arm and all seven seeded arms, and each cluster is counted once per set. (a) Full versus seeded arm under the same homology annotation (II versus III), counted as distinct Swiss-Prot accessions (**Table 1, S1**) and summed over the seven classes, (b) Full arm with DIAMOND (I) versus REFinder seeded arm with REBEAN (IV), counted as enzyme clusters because REBEAN enzymes need not have a Swiss-Prot hit. (c) Known and novel enzyme clusters in I, in IV, and in IV by EC class; a cluster is novel if no member has a Swiss-Prot hit at 30% identity or above. Boxes show the median and interquartile range (IQR), whiskers extend to the most extreme value within 1.5 × IQR, and points are individual samples; y axes in log scale.

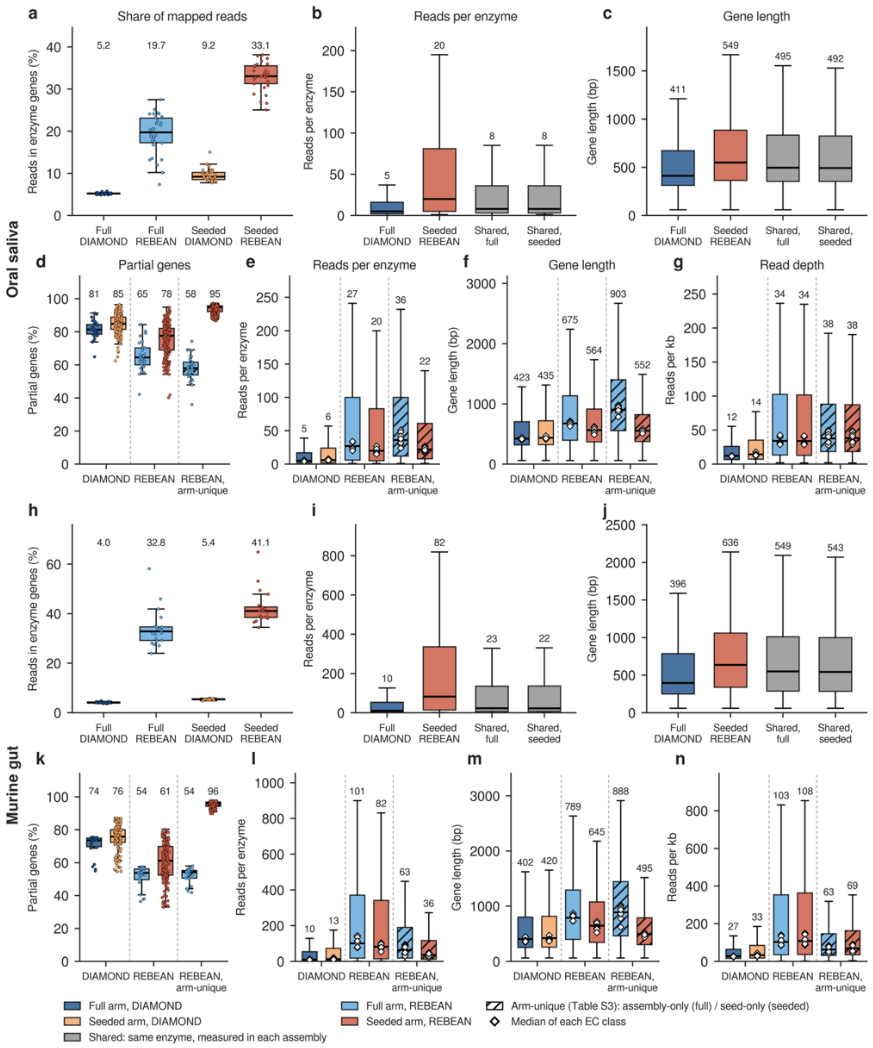


**Figure S4: Read-to-gene map, gene length and completeness by full and seeded arms.** Panels a–g show oral saliva (PRJNA396840, n = 30) and h–n murine gut (PRJNA504846, n = 20) metagenomes. Enzymes are within-sample MMseqs2 clusters of predicted proteins (90% identity, 80% coverage), compared class-matched: EC-N enzymes of the full arm against those seeded arm, pooled over EC1–7. A protein is a DIAMOND enzyme if its best Swiss-Prot hit (blastp, $E \leq 10^{-5}$) carries a class-N EC number, and a REBEAN enzyme if a mapped read reaches the class-N 90%-precision threshold. Read counts are of each sample's fastq screened reads aligned to the respective assembly (BBMap, ≥ 90% identity) whose primary alignment overlaps the gene. (a, h), Percentage of each assembly's mapped reads that lie in enzyme genes, per sample, EC. (b, c, i, j), Reads per enzyme and gene length for full-assembly DIAMOND enzymes, seeded-assembly REBEAN enzymes and the enzymes shared by both, each shared enzyme shown as measured in either assembly; a-c and h-j use one representative gene per enzyme. (d, k), Percentage of partial Prodigal gene calls per assembly for all DIAMOND and REBEAN enzymes of each arm and for arm-unique REBEAN enzymes (hatched), which are assembly-only (full arm) or seed-only (seeded arm). (e-g, l-n), Reads per enzyme, gene length and read depth (reads per kb) for the same sets, with reads and length summed over each arm's annotated genes within a cluster. Colors denote arm and annotation: dark blue, full arm, DIAMOND; light orange, seeded arm, DIAMOND; light blue, full arm, REBEAN; red, seeded arm, REBEAN; grey, shared. Boxes show the median and interquartile range (IQR); whiskers extend to the most extreme value within 1.5 × IQR; outliers are not shown. Points are samples (a, h) or assemblies (d, k). Numbers and white diamonds denote the median of each EC class.

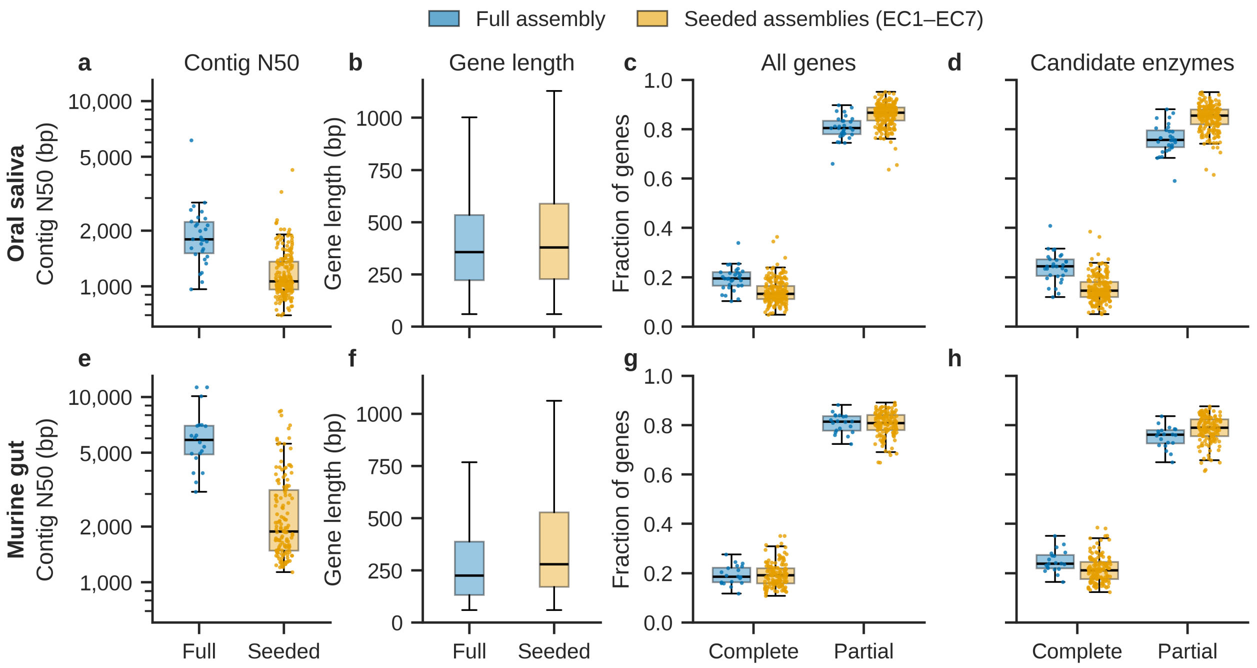


**Figure S5: Assembled contig and gene length of full and REBEAN-seeded assemblies.** Panels a-d show the oral saliva metagenomes (30 samples) and panels e–h the murine gut metagenomes (20 samples). In blue: the full assembly of all quality-filtered reads and in orange: the seven EC-seeded assemblies (EC1–EC7) of each sample (n = 210 oral, 140 murine). (a, e) Contig N50, computed over contigs of at least 500 bp; (b, f) Length of every Prodigal gene call, pooled across the assemblies of each arm: 9.6 million (full) and 18.7 million (seeded) genes in the oral set; 22.5 million and 49.9 million in the murine set. (c, g) Fraction of Prodigal gene calls per assembly that are complete or partial. (d, h) As in c and g, restricted to candidate enzyme genes. These are genes with at least one read that REBEAN classifies as enzymatic, or whose best Swiss-Prot hit (DIAMOND blastp, $E \leq 10^{-5}$) carries an EC number. They are the genes clustered to define distinct enzymes in Tables 2, S3 and S4. Boxes show the median and interquartile range. Whiskers extend to the most extreme value within 1.5 × the interquartile range, and outliers are not drawn. Points are individual assemblies. In b and f, quartiles are computed over all genes.
